# Dual Value System and its assessment scheme for understanding and valuing ecosystem services

**DOI:** 10.1101/628578

**Authors:** Haile Yang, Jiakuan Chen

## Abstract

Valuing ecosystem services (ES) is helpful for effective ES management. However, there are many limitations in traditional ES valuation approaches, including theoretical challenges and practical difficulties. To overcome these limitations, we proposed a dual value system (DVS). And then, we presented a case study of valuing the water provision in Zhujiang River Basin (Pearl River Basin) based on DVS. DVS follows the axioms that (1) human life would end if we lose any of vital ES which is indispensable to human being’s survival (such as oxygen, freshwater) and (2) ES cannot provide any value to people without human activities. Correspondingly, DVS includes two types of value: the output support value (OSV) of a vital ES refers to the total value produced by human being’s economic and social activities (TVPH) supported by the ES consumption; the optional capacity value (OCV) of a vital ES refers to the optional capacity of supporting TVPH provided by total ES volume. The OCV provided by a vital ES is calculated by using the product of multiplying the OSV (TVPH) by the freedom of choosing the consumption from the total volume of this ES, valued in non-monetary units. Based on DVS, the OSV and OCV of water provision in Zhujiang River Basin were analyzed in river basin scale and sub-basin scale, and the values variation of water provision from 2006 to 2015 was analyzed in sub-basin scale. And then, based on this case study, we discussed the new insights into ES provided by DVS. Results proved that DVS and its assessment scheme overcame the limitations on current ES valuation approaches and provided an innovative quantitative framework to understand and value ES which will help to make good decisions in ES management.

## 1. Introduction

Ecosystems provide a range of services that are fundamentally important for human wellbeing, health, livelihoods, and survival (Costanza et al., 1997; Millennium Ecosystem Assessment, 2005; TEEB Synthesis, 2010). To unravel the complex socio-ecological relationships and explicate how human decisions would affect ecosystem services (ES), the values of ES are used to express these changes which allow for their incorporation in public decision-making processes (Farley and Costanza, 2010; Pascual et al., 2010; Costanza et al., 2017).

The valuation of ES could be monetary, non-monetary, or mixes of monetary and non-monetary (Figure 1). In monetary ES valuation studies, there are two approaches (Jiang, 2017): (1) unit value based approach (i.e. benefit transfer) (Costanza et al., 1997; de Groot et al., 2012; Costanza et al., 2014; Kubiszewski et al., 2017) and (2) primary data based approach (Ouyang et al., 1999; Guo et al., 2001; Dai et al., 2016) (Figure 1). In non-monetary ES valuation studies, there are some biophysical approaches (Koellner and Geyer, 2013; Coscieme et al., 2014; Mancini et al., 2018), such as Emergy analysis, Ecological Footprint accounting; and some pluralistic approaches (Pascual et al., 2017; Martin and Mazzotta, 2018; Folkersen, 2018), such as the pluralistic valuation approach provided by Intergovernmental Science-Policy Platform on Biodiversity and Ecosystem Services (IPBES). IPBES considers that the ES (redefined as nature’s contributions to people) and a good quality of life are interdependent, and then processes biophysical, sociocultural, economic, health and holistic valuations (Pascual et al., 2017) (Figure 1). The approaches of valuing ES using mixes of monetary and non-monetary units are diverse (Hayha et al., 2015; Kenter et al., 2016) (Figure 1). In the case provided by Hayha et al. (2015), the ES are assessed in biophysical and monetary units (Hayha et al., 2015). Kenter et al. (2016) integrate deliberative monetary valuation, storytelling, subjective wellbeing and psychometric approaches to comprehensively elicit cultural ecosystem services values (Kenter et al., 2016).

**Figure 1.**
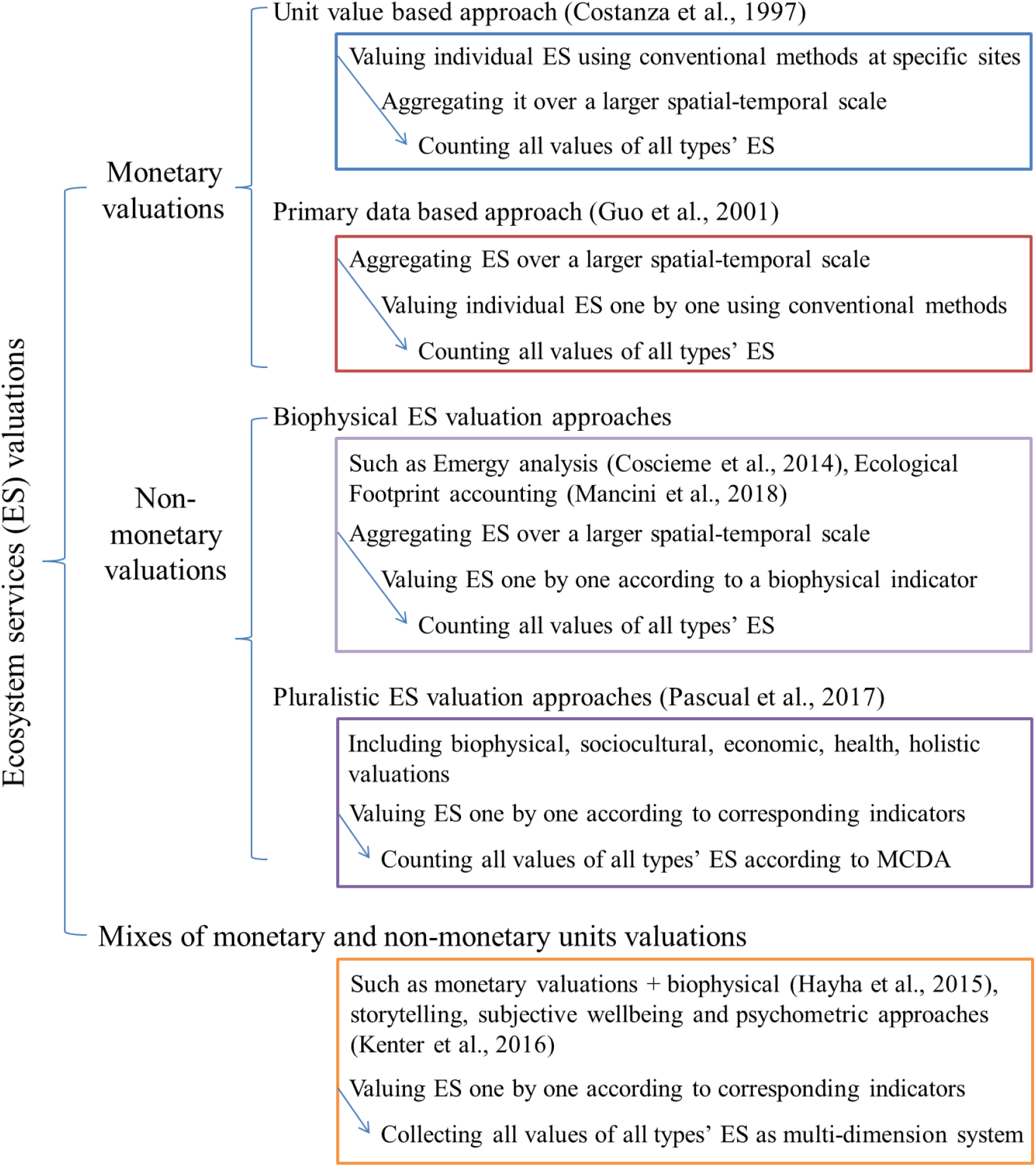
Types of ecosystem services valuation

There are a series of limitations on current valuation methods. Theoretically, there are two challenges in current valuation methods (Costanza et al., 2017): (1) imperfect information, for example individuals might assign no value on an ES if they do not know the role that the service is playing in their wellbeing (Norton et al., 1998; van Ree et al., 2017); (2) the difficulties of accurately measuring the system’s functioning so as to correctly quantify a given service value derived from that system (Boumans et al., 2002; Barbier et al., 2008; Koch et al., 2009). These two challenges lead to two main problems in valuing the overall ES value of a region (or the Earth): the miss-counting or double-counting of ES types, and the underestimate or overestimate of individual ES. And then, according to current aggregation methods, these two problems lead to a more serious underestimate or overestimate on the overall ES value of a region (or the Earth).

Practically, there are two difficulties in current valuation methods: (1) most of the current monetary ES valuation approaches are based on the conventional valuation techniques which are diverse and limited (Pascual et al., 2010; Folkersen, 2018); (2) most of the current non-monetary ES valuation approaches use multi-indicators (Pascual et al., 2017; Martin and Mazzotta, 2018), which make valuation approaches complex and costly. To solve the amplified underestimate or overestimate on the overall ES value of a region (or the Earth) caused by theoretical challenges and current aggregation approaches, and to avoid the practical difficulties in valuing ES, we need to propose a new assessment framework.

In this work, we firstly proposed a dual value system (DVS), including two new concepts: output support value (OSV) and optional capacity value (OCV). And then, we presented a case study of analyzing the OSV and OCV of water provision in Zhujiang River Basin (Pearl River Basin) to demonstrate the application of DVS. In this case study, to explore the new insights of DVS into ES and the innovative assessment scheme of DVS, we (1) analyzed the OSV and OCV of water provision at two spatial scales: river basin scale and sub-basin scale, (2) analyzed the OCV of the passing-by water among hydrologic units at the sub-basin scale, (3) analyzed the OSV and OCV variation of the water provision from 2006 to 2015.

## 2. Methods and materials

### 2.1 Methods details: the DVS for valuing ES

Chaisson (2002) has argued that the global value of ES is infinite, as we human beings could not survive if we lost any vital ES which is indispensable to human being’s survival (such as oxygen, freshwater). The truth indicated by him shows that the total value produced by human being’s economic and social activities (TVPH) relies on each vital ES. Costanza et al. (2014) have indicated that ecosystems could not provide any values to people without the presence of people (human capital), their communities (social capital), and their built environment (built capital). In other words, the value of each ES depends on the level of socioeconomic development. Here we get two inferences based on these two axioms.

The first inference is that the consumption of vital ES supports the TVPH. To describe this inference, we proposed a new concept which is termed as “Output Support Value (OSV)”. OSV of a vital ES refers to the TVPH supported by the ES consumption, which shows the benefits for human wellbeing derived from consuming a definite volume of ES, directly or indirectly.

The second inference is that the total volume of a vital ES provides the freedom of consuming corresponding ES in maintaining the human being’s production and survival. To describe this inference, we proposed a new concept which is termed as “Optional Capacity Value (OCV)”. OCV of a vital ES refers to the optional capacity of supporting the OSV (i.e. TVPH) provided by the total volume of ES, which shows the benefits for human wellbeing derived from having the option of using, directly or indirectly. OCV is described as the product of multiplying the OSV (i.e. TVPH) by the freedom of choosing the consumption from the total volume of ES.

Based on these two inferences and two new concepts, we constructed a brand new assessment scheme for valuing ES – “dual value system (DVS)”, in which OSV was valued in monetary units and OCV was valued in non-monetary units.

For valuing ES, we need first to classify ES in an appropriate way. The classification of ES mainly has two approaches (Hayha and Franzese, 2014). The first approach is the classification mainly based on four categories: (1) provisioning, (2) regulating, (3) supporting, and (4) cultural services, which is the most commonly used classification (Millennium Ecosystem Assessment, 2005; TEEB Synthesis, 2010; CICES – Haines-Young and Potschin, 2013). And the second approach is the classification mainly based on spatial characteristics of ES (Costanza, 2008; Fisher et al., 2009). To satisfy the needs of valuing ES in DVS, we proposed a classification of ES based on indispensability and spatial characteristics of ES (Table 1) (Figure 2), following the second approach.

**Figure 2.**
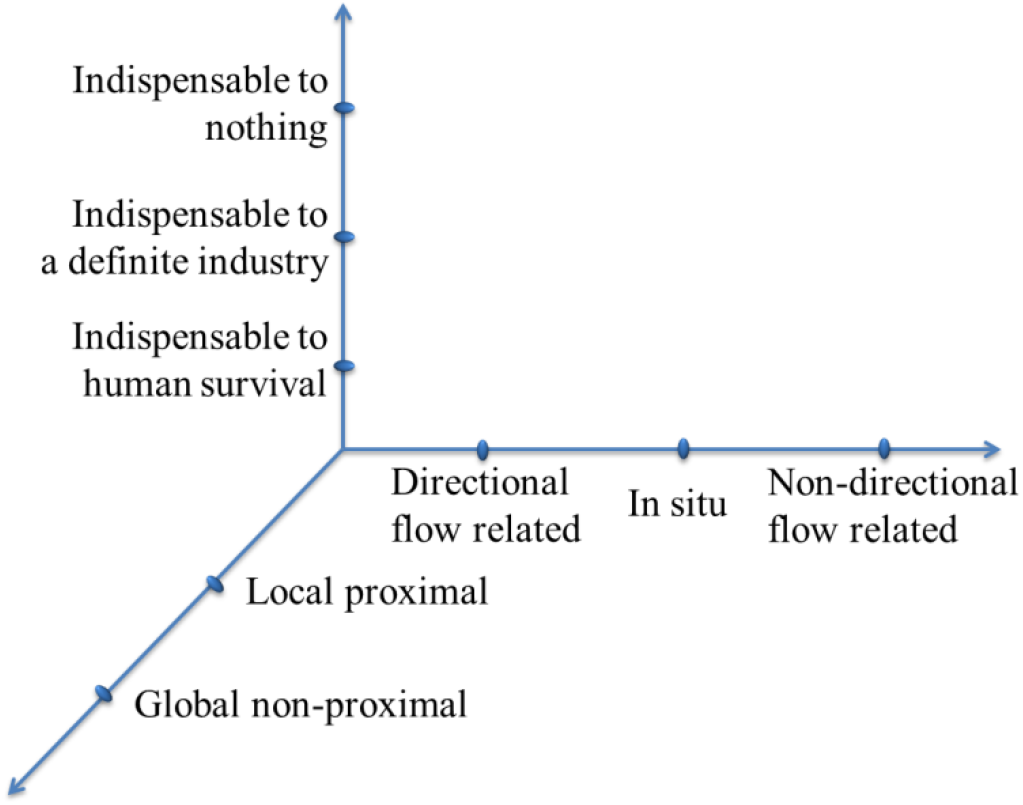
Three-dimensional schematic diagram for the classification of ecosystem services based on indispensability and spatial characteristics. For example, oxygen is indispensable to human survival, global non-proximal, non-directional flow related; fresh water is indispensable to human survival, local proximal, directional flow related.

**Table 1.** Classification of ecosystem services based on indispensability and spatial characteristics, modified from Hayha and Franzese (2014)

| Classification | Example |
| --- | --- |
| Indispensable to human survival | Basic services for human survive; e.g. fresh water, oxygen |
| Indispensable to a definite industry | Basic services for an industry; e.g. honeybee |
| Indispensable to nothing | Services that human cannot use; e.g. hydrothermal vent ecosystem |
| Global non-proximal | Does not depend on proximity; e.g., carbon sequestration |
| Local proximal | Depends on proximity; e.g., disturbance regulation |
| Directional flow related | Flow from point of production to point of use; e.g., water supply |
| In situ | Point of use; e.g., soil formation |
| Non-directional flow related | Flow from point of production to adjacent area of use; e.g., climate regulation |

In our classification of ES (Figure 2), indispensability means that if we lose any vital ES (indispensable to human survival, such as oxygen, freshwater), human life would end; if we lose crucial ES (indispensable to definite industries, such as honeybee), all industries based on this ES would be vanished. In DVS, we only value the currently usable ES. And an individual ES is valued only based on current socioeconomic status.

Regarding the spatial characteristics, there are ES that are not dependent on the specific location (e.g., global carbon sequestration, global oxygen production), ES that are instead dependent on their spatial distribution in relation to human presence (e.g., waste treatment and storm protection), ES in which the direction of the flow from upstream to downstream matters (e.g., water supply and sediment transport), ES in which the flow without definite direction matters (e.g., fresh air supply and climate regulation), and ES in which no flow matters (e.g., soil formation).

Following our classification of ES (Figure 2), OSV of the ES which is indispensable to human survival could be valued using TVPH; OSV of the ES which is indispensable to definite industries could be valued using the TVPH based on these industries (Table 2). OCV of an ES is calculated as the product of multiplying its OSV by the freedom of choosing the ES consumption from the total volume of ES. The freedom is evaluated by the average uncertainty which could be described by log base 2 which indicates the uncertainty in a binary decision, and is valued in bits (Ulanowicz, 1986) (Table 2).

**Table 2.** The Output Support Value (OSV) and Optional Capacity Value (OCV) of an ecosystem service (ES)

| Ecosystem service (ES) | Output Support Value (OSV) | Optional Capacity Value (OCV) |
| --- | --- | --- |
| Indispensable to human survival | $OSV = TVPH$ | $OCV = OSV * \log_2 \frac{ES_t}{ES_c}$ |
| Indispensable to a definite industry | $OSV = TVPH$ based on corresponding industry | $OCV = OSV * \log_2 \frac{ES_t}{ES_c}$ |

In the equation, $ES_t$ denotes the total volume of an ecosystem service in a region; $ES_c$ denotes the consumed volume of an ecosystem service in maintaining the production and survival, or in supporting a definite industry; TVPH denotes the total value produced by human being's economic and social activities.

Regarding the spatial characteristics, we take regions as the accounting units. In another word, each accounting unit provides a definite ES value. As the global non-proximal ES supports the global human activities, it is valued by global TVPH. As the local proximal ES only supports the local human activities, it is valued by local TVPH. For the ES with (directional or non-directional) flow, both the local ES in a region and the imported ES come from other regions (accounting units) support human activities in this region. Following the principle of local priority, the value of imported ES is assigned as the value that the value of total ES (sum of the local ES and the imported ES) minus the value of local ES (Eq. 1, 2, 3, 4). For the ES in which no flow matters, its value is the value of local ES.

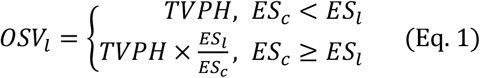

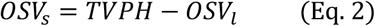

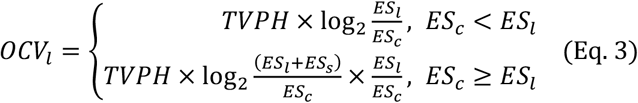

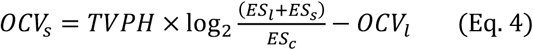

In the equations, *OSV_ι_* denotes the OSV of local ecosystem service (ES) in a region; *OSV_s_* denotes the OSV of ES spatial inflow in a region; *OCF_ι_* denotes the OCV of local ES in a region; *ES_c_* denotes the consumed volume of ES in a region; TVPH denotes the total value produced by human being’s economic and social activities.

In the framework of DVS, all types of ES could be valued parallelly. The overall value of ES in a region (or the Earth) takes the maximum one from the values of all types’ ES (Figure 3).

**Figure 3.**
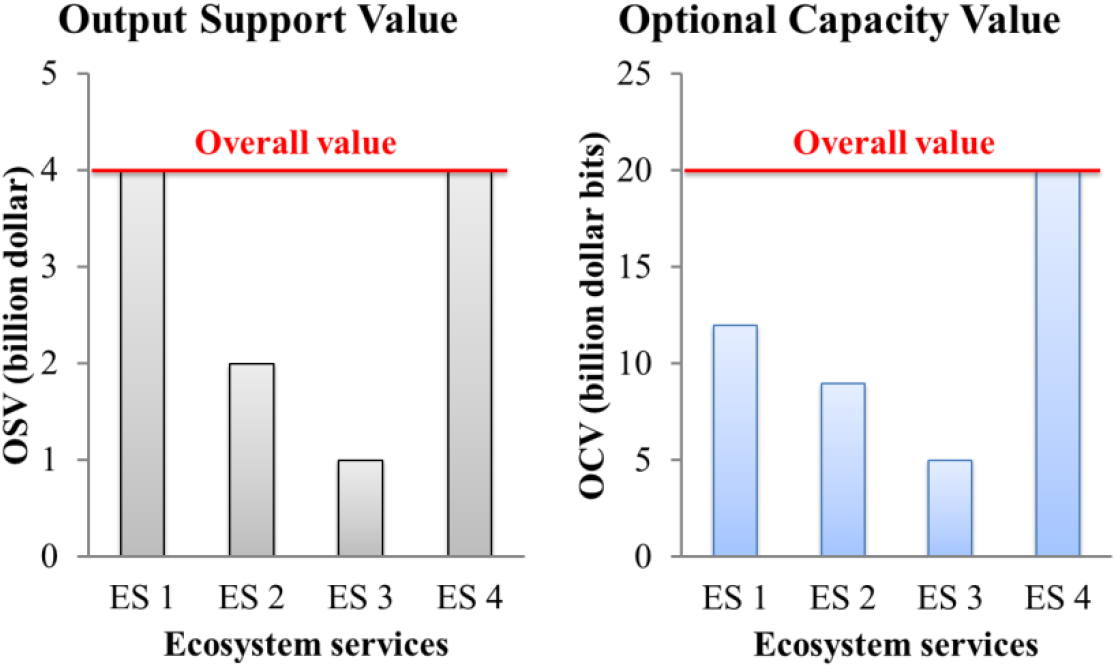
Schematic representations of valuing the overall value of multi-types’ ecosystem services (ES). ES 1 represents an ecosystem service which is indispensable to human survival; and ES 4 represents another ecosystem service which is indispensable to human survival; ES 2 represents an ecosystem service which is indispensable to a definite industry; ES 3 represents another ecosystem service which is indispensable to another definite industry. The values in these figures are all fictitious.

### 2.2 Study area: the Zhujiang River Basin

In DVS framework, to value ES, one needs three sets of data with spatial-temporal consistency: total ES volume, ES consumption and TVPH. Moreover, to discuss the new insights of DVS into ES value, such as the scale-specificity of DVS in valuing ES and the efficiency of DVS in valuing the ES fluxes, the case study area should be nested divided and each subdivision has the same data sets, the data on ES fluxes among subdivisions should be available. The water provision in Zhujiang River Basin meets these conditions.

Here, we presented a case study of valuing the water provision in Zhujiang River Basin based on DVS framework. As water resources are indispensable to human survival and local proximal, following DVS, we analyze the value of water provision based on total water resources volume, water consumption and TVPH. Moreover, as water resources are directional flows, we delineate the donors and recipients of water resources fluxes (passing-by water), and then value each fluxes based on the volume of passing-by water, water consumption and TVPH of recipients. In Zhujiang River Basin, the Zhujiang River Water Resources Bulletin (1) provides the total water resources volume, water consumption and gross domestic product (GDP, here indicates TVPH) of the overall river basin, (2) divides the Zhujiang River Basin into seven hydrologic units: the Nanpan-Beipan River, the Hong-Liu River, the Yujiang River, the Xijiang River Lower Reach, the Beijiang River, the Dongjiang River and the Zhujiang Delta (Figure 4), (3) provides the total water resources volume, water consumption and GDP of each hydrologic units and (4) provides the data for calculating the volume of passing-by water among hydrologic units, and the water consumption and GDP of recipients (PRWRC, 2017).

**Figure 4.**
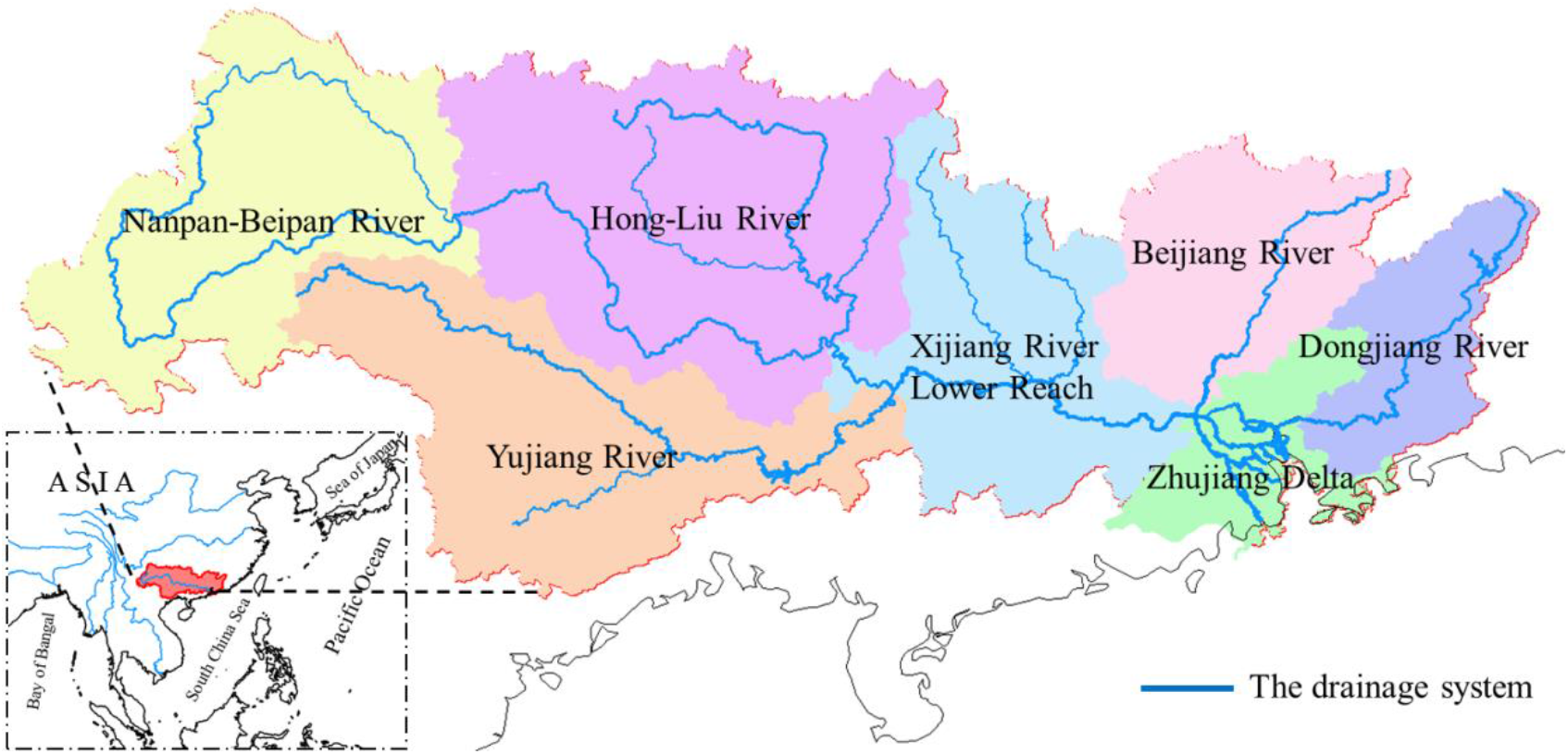
Zhujiang River Basin (Pearl River Basin) and its seven hydrologic units.

As a case study for demonstrating the application of DVS, in valuing the water provision in Zhujiang River Basin, we only consider the volume of water resources rather than its environment and temporal distribution for brevity. All data that we use are available from the Zhujiang River Water Resources Bulletin provided by the Pearl River Water Resources Commission of the Ministry of Water Resources (PRWRC) (http://www.pearlwater.gov.cn/xxcx/szygg/).

## 3. Results

### 3.1 The value of the water provision at river basin scale

Firstly, we valued the water provision at river basin scale by taking the Zhujiang River Basin as the accounting unit. In 2015, as the total volume of water resources in Zhujiang River Basin is 410.04 billion m^3^, the water usage 60.43 billion m^3^ and the GDP 8.5105 trillion yuan (PRWRC, 2017), the OSV of the water provision in Zhujiang River Basin is 8.5105 trillion yuan which indicates that the water usage of 60.43 billion m3 supports the GDP of 8.5105 trillion yuan; the OCV of the water provision in Zhujiang River Basin is 23.5096 trillion yuan bits which indicates that the total water provision of 410.04 billion m^3^ provides the optional capacity for supporting the TVPH (measured by GDP) as much as 23.5096 trillion yuan bits (Table 3).

**Table 3.** The Output Support Value (OSV) and Optional Capacity Value (OCV) of water provision provided and experienced by Zhujiang River Basin at river basin scale in 2015

|  | Total volume<br>(e9 m<sup>3</sup>) | Usage (e9<br>m<sup>3</sup>) | GDP (e12<br>yuan) | OSV (e12<br>yuan) | OCV (e12<br>yuan bits) |
| --- | --- | --- | --- | --- | --- |
| Zhujiang<br>River Basin | 410.04 | 60.43 | 8.5105 | 8.5105 | 23.5096 |

### 3.2 The value of the water provision at sub-basin scale

Secondly, we valued the water provision at sub-basin scale by taking seven major hydrologic units of the Zhujiang River Basin as the accounting units. For each hydrologic unit, both the passing-by water and the local water yield provide the optional capacity for supporting the TVPH (measured by GDP). Following the principle of local priority, in 2015, the Hong-Liu River, Xijiang River Lower Reach and Zhujiang Delta respectively receive passing-by water 37.40 billion m^3^, 200.28 billion m^3^ and 360.20 billion m^3^ which provide OCV 0.1935 trillion yuan bits, 1.0332 trillion yuan bits and 18.7463 trillion yuan bits, respectively (Table 4).

**Table 4.** The Output Support Value (OSV) and Optional Capacity Value (OCV) of water provision experienced by each hydrologic unit of Zhujiang River Basin at sub-basin scale in 2015

|  | Total volume<br>(e9 m<sup>3</sup>) | Usage (e9<br>m<sup>3</sup>) | GDP (e12<br>yuan) | OSV (e12<br>yuan) | OCV (e12<br>yuan bits) |
| --- | --- | --- | --- | --- | --- |
| Nanpan-Beipan River | 40.00 | 4.77 | 0.5817 | 0.5817 | 1.7846 |
| Hong-Liu River | 126.37 | 9.78 | 0.5175 | 0.5175 | 1.9103 |
| **Passing-by water** | **37.40** |  |  |  | **0.1935** |
| Yujiang River | 44.14 | 7.86 | 0.5240 | 0.5240 | 1.3045 |
| Xijiang River Lower<br>Reach | 79.92 | 10.79 | 0.5709 | 0.5709 | 1.6492 |
| **Passing-by water** | **200.28** |  |  |  | **1.0332** |
| Beijiang River | 61.68 | 5.26 | 0.3757 | 0.3757 | 1.3344 |
| Dongjiang River | 27.40 | 4.64 | 0.8436 | 0.8436 | 2.1614 |
| Zhujiang Delta | 30.53 | 17.33 | 5.0971 | 5.0971 | 4.1641 |
| **Passing-by water** | **360.20** |  |  |  | **18.7463** |
| **Sum** | **1007.92** | **60.43** | **8.5105** | **8.5105** | **34.2816** |

Distributing the OCV of the passing-by water into every hydrologic unit which exports water resources, we calculated the OCV of the water provision provided by each hydrologic unit, including local service and output service (Table 5). The result showed the OCV of the point-to-point water provision fluxes (Table 5).

**Table 5.** The Optional Capacity Value (OCV) of water provision provided by each hydrologic unit of Zhujiang River Basin in 2015

|  | OSV (e12 yuan bits) |  |  |  | Sum |
| --- | --- | --- | --- | --- | --- |
|  | Local service | Output to Hong-Liu River | Output to Xijiang River Lower Reach | Output to Zhujiang Delta |  |
| Nanpan-Beipan River | 1.7846 | 0.1935 | 0.1929 | 1.9465 | 4.1176 |
| Hong-Liu River | 1.9103 |  | 0.6311 | 6.3671 | 8.9085 |
| Yujiang River | 1.3045 |  | 0.2091 | 2.1099 | 3.6235 |
| Xijiang River Lower Reach | 1.6492 |  |  | 3.8976 | 5.5468 |
| Beijiang River | 1.3344 |  |  | 3.0857 | 4.4201 |
| Dongjiang River | 2.1614 |  |  | 1.3396 | 3.5010 |
| Zhujiang Delta | 4.1641 |  |  |  | 4.1641 |

### 3.3 The value variation of the water provision from 2006 to 2015 at sub-basin scale

Thirdly, we compared the OSV and OCV of the water provision provided by each hydrologic unit in 2006 and 2015. Results showed that the OCV of the water provision of both the local and the exported services provided by each hydrologic unit increase with the economic growth (Table 6).

**Table 6.** The Output Support Value (OSV) and Optional Capacity Value (OCV) of local and passing-by water provision provided by each hydrologic unit of Zhujiang River Basin in 2006 and 2015

|  | GDP<br>(e12 yuan) |  | Water resources<br>(e9 m<sup>3</sup>) |  | Water usage<br>(e9 m<sup>3</sup>) |  | OSV (e12 yuan) |  | OCV (e12 yuan bits) |  |  |  |  |  |
| --- | --- | --- | --- | --- | --- | --- | --- | --- | --- | --- | --- | --- | --- | --- |
|  |  |  |  |  |  |  |  |  | Local service |  | Output service |  | Sum |  |
|  | 2006 | 2015 | 2006 | 2015 | 2006 | 2015 | 2006 | 2015 | 2006 | 2015 | 2006 | 2015 | 2006 | 2015 |
| Nanpan-Beipan River | 0.1577 | 0.5817 | 29.66 | 40.00 | 4.62 | 4.77 | 0.1577 | 0.5817 | 0.4231 | 1.7846 | 0.6269 | 2.3329 | 1.0500 | 4.1176 |
| Hong-Liu River | 0.1481 | 0.5175 | 87.35 | 126.37 | 10.34 | 9.78 | 0.1481 | 0.5175 | 0.4560 | 1.9103 | 1.7373 | 6.9982 | 2.1934 | 8.9085 |
| Yujiang River | 0.1442 | 0.5240 | 36.95 | 44.14 | 7.29 | 7.86 | 0.1442 | 0.5240 | 0.3374 | 1.3045 | 0.6981 | 2.3190 | 1.0355 | 3.6235 |
| Xijiang River Lower Reach | 0.1765 | 0.5709 | 70.74 | 79.92 | 11.82 | 10.79 | 0.1765 | 0.5709 | 0.4555 | 1.6492 | 1.2427 | 3.8976 | 1.6982 | 5.5468 |
| Beijiang River | 0.1055 | 0.3757 | 64.20 | 61.68 | 5.36 | 5.26 | 0.1055 | 0.3757 | 0.3780 | 1.3344 | 1.1734 | 3.0857 | 1.5514 | 4.4201 |
| Dongjiang River | 0.2837 | 0.8436 | 38.32 | 27.40 | 4.20 | 4.64 | 0.2837 | 0.8436 | 0.9051 | 2.1614 | 0.6961 | 1.3396 | 1.6011 | 3.5010 |
| Zhujiang Delta | 1.7566 | 5.0971 | 34.26 | 30.53 | 19.50 | 17.33 | 1.7566 | 5.0971 | 1.4286 | 4.1641 |  |  | 1.4286 | 4.1641 |
| Sum | 2.7723 | 8.5105 | 361.49 | 410.04 | 63.13 | 60.43 | 2.7723 | 8.5105 | 4.3836 | 14.3085 | 6.1745 | 19.9731 | 10.5581 | 34.2816 |

## 4. Discussion

### 4.1 New insights of DVS

DVS provides a new perspective to understand the value of ES. In 2015, in Zhujiang River Basin at the river basin scale, the OSV of the water provision (8.5105 trillion yuan) means that a definite volume water usage (60.43 billion m^3^) supports a definite TVPH (8.5105 trillion yuan, measured by GDP) (Table 3). The OCV of the water provision (23.5096 trillion yuan bits) means that the volume of total water provision (410.04 billion m^3^) provides the optional capacity for supporting the TVPH (23.5096 trillion yuan bits, measured by GDP) (Table 3). DVS makes the importance and value of ES easy be understood through direct value (OSV) and indirect value (OCV) in the perspective different from traditional exchange value.

DVS provides new insight into the relationship between the ES value and the social and economic development (Table 2). From 2006 to 2015, the OSV and OCV of the water provision in seven accounting units of Zhujiang River Basin increase with their TVPH (here measured by GDP) (Table 6). This is consistency with the point in previous researches (Costanza et al., 2014). Conclusively, the social and economic development depends on the ES and the ES value increases with the social and economic development. Based on this, we would not simply make the social and economic development against with the ecological conservation.

DVS provides new insight into how to understand and quantify the value of ES spatial subsidies (i.e. ES fluxes from one region to another region). The spatial subsidies of ES increase (1) the ES OCV experienced by the recipient accounting units and (2) the ES OCV provided by the donor accounting units, and then (3) the social and economic development of recipient accounting units increase the ES OCV provided by the donor accounting units. In Zhujiang River Basin, according to the passing-by water, (1) the recipients receive more OCV (Table 4), (2) the donors provide more OCV (Table 5), and (3) the economic growths in recipients promote the increase of the OCV provided by donors (Table 6). Based on this framework, we could evaluate the ES value variation in different scenarios of ecological conservation and natural resources exploitation in a region, and then optimize the plans for ecological conservation and natural resources exploitation.

DVS provides a new method to quantify payments for ecosystem services (PES). ES spatial subsidies provide OCV to recipients (accounting unit), then recipients should provide corresponding ecological compensation to donors (Yang et al., 2019). The compensation standard could reference to the OCV of ES spatial subsidies. In 2015, the hydrologic unit of Nanpan-Beipan River provides the OCV of the water provision 1.7846 trillion yuan bits, 0.1935 trillion yuan bits, 0.1929 trillion yuan bits, 1.9465 trillion yuan bits to the Nanpan-Beipan River, Hong-Liu River, Xijiang River Lower Reach and Zhujiang Delta respectively (Table 5). Then, if a water resources protection program in Nanpan-Beipan River costs 4.1176 billion yuan, the Nanpan-Beipan River, Hong-Liu River, Xijiang River Lower Reach and Zhujiang Delta should pay 1.7846 billion yuan, 0.1935 billion yuan, 0.1929 billion yuan, 1.9465 billion yuan respectively.

DVS provides a new framework for ES management. Following DVS, one could (1) identify the features of an individual ES, (2) identify the accounting units appropriately, (3) collect corresponding data sets, and then (4) calculate the OSV and OCV of this ES (Table 2). Based on the ES OCV, one could discuss the ES management in each region and the PES among regions (Yang et al., 2019). If one values a set of ES in a region with limited space and resources, one could estimate the overall ES value in this region following DVS (Figure 3), and then discriminate between different services to decide what should be managed and what should be traded to satisfy human needs/ wants for human wellbeing. In this case study, we valued the water provision in Zhujiang River Basin using the total water provision volumes, water consumptions and TVPH (indicated by GDP) at basin scale and sub-basin scale respectively. This method could be used in any other river basins and regions.

In DVS, there are intrinsic consistency among the OSV at different scales and intrinsic incommensurability among the OCV at different scales. OSV of an ES means the TVPH supported by the consumed ES, and the TVPH and consumed ES in a certain space-time condition are constant, which is independent of spatial and temporal scales, so OSV are intrinsically consistent at different scales. In contrast, OCV of an ES means the optional capacity of supporting the TVPH provided by the total ES, and the spatial subsidies of an ES among accounting units increase the total volume of ES in every accounting unit, their OCV are intrinsically incommensurable at different scales. Moreover, the spatial distributions of the TVPH, ES and ES consumption are mismatched, and OCV of an ES is calculated by these three indicators non-linearly (Table 2), so their OCV (even only considering the local services) are intrinsically incommensurable at different scales.

### 4.2 Advantages of DVS

DVS provides a simple and general assessment scheme for valuing ES. Firstly, DVS avoids the practical difficulties in traditional ES valuation approaches, such as the difficulty of re-estimating a large amount of unit values and the difficulty of accurately valuing each individual ES. Secondly, the misestimate of ES value in DVS is traceable and easy-to-correct, although DVS does not completely avoid the underestimate or overestimate on individual ES led by insufficient information and misunderstanding on an ES. Thirdly, DVS provides a general approach and a general unit to value ES, avoids the difficulties for transforming and aggregating incommensurable data which come from the values of different type’s ES. Fourthly, DVS avoids the amplification of underestimate or overestimate on the overall ES value of a region (or the Earth), although DVS do not completely avoid the miss-counting or double-counting of ES types led by insufficient information.

DVS avoids the practical difficulties in traditional ES valuation approaches. The value of ES depends on the level of socioeconomic development (Costanza et al., 2014). So, following the unit value based approach, one should value the 2011 ES using the 2011 unit values (Costanza et al., 2014), value the 2050 ES using the 2050 unit values (Kubiszewski et al., 2017), rather than value the 2011 ES using the 1997 unit values (Costanza et al., 2014), value the 2050 ES using the 2011 unit values (Kubiszewski et al., 2017). However, re-estimating all unit values is a heavy work, especially, considering the non-linearity in ecosystem services (Barbier et al., 2008; Koch et al., 2009) and the spatial heterogeneity of unit values (de Groot et al., 2012; Crossman et al., 2013; Drakou et al., 2015). In the primary data based approach, the difficulty for accurately valuing each individual ES is remained (Guo et al., 2001; Dai et al., 2016), although there are some technique innovations in recent years (Coscieme et al., 2014). In DVS, the OSV of an ES is determined by the TVPH and ES consumption; the OCV of an ES is determined by total ES volume, ES consumption and OSV (Table 2). In this case study, the TVPH was indicated by GDP which is the only indicator of indicating the TVPH in all current social and economic indexes. If someone gets another indicator which could indicate the TVPH more accurately, he could use that indicator to replace GDP in his study. This replacement has no impact on the assessment scheme of DVS. In this work, the freedom was evaluated by the average uncertainty of selecting ES consumption from the total volume of this ES. The average uncertainty was described by log base 2 which indicated the uncertainty in a binary decision (Ulanowicz, 1986). If someone believes that another indicator could indicate the average uncertainty more appropriately, he could use that one to replace log base 2 in his study. This replacement does not change the assessment scheme of DVS. So, DVS reflects the level of economic development and avoids the difficulties in unit value based approach and primary data based approach.

DVS makes the misestimate of ES value traceable and easy-to-correct. In traditional approaches, such as direct market valuation methods, revealed preference methods and stated preference methods, the valuation of different types of ES depends on different rules and their underestimate and overestimate are difficult to identify and correct (Pascual et al., 2010). The ES valuation method in DVS (Table 2) makes the underestimate or overestimates of individual ES value to be traceable and easy-to-correct, although DVS does not completely avoid the underestimate or overestimate on individual ES led by insufficient information and misunderstanding on an ES.

DVS provides a general approach and a general unit to value ES. There is a great difficulty in aggregating the non-monetary values of ES because the non-monetary ES values could be qualitative or quantitative, and are valued by diverse units/ indicators for different ES. For resolving this problem, someone tries the Multi-Criteria Decision Analysis (MCDA) so as to transform the incommensurable data (i.e., monetary and non-monetary values) into non-monetary, dimensionless values, and then to mathematically incorporate non-monetary features into the aggregation (Martin and Mazzotta, 2018). However, one needs to provide some assumptions if he/ she wants to transform a set of incommensurable data and aggregate them. Different researchers provide different assumptions, and their aggregating results are sensitive to their assumptions (Martin and Mazzotta, 2018). Moreover, the transforming and aggregating approaches are very complex. In DVS, we calculate the OCV of all types’ ES using a general approach and a general unit. So, DVS avoids the difficulties for transforming and aggregating incommensurable data which come from the values of different type’s ES.

DVS avoids the amplification of underestimate or overestimate on the overall ES value of a region (or the Earth). The common problem of valuing the total economic value of ES in a region (or the Earth) in both the unit value based approach and the primary data based approach is the amplified underestimate or overestimate caused by the miss-counting or double-counting of ES types (Fu et al., 2011; Stoeckl et al., 2014; van Ree et al., 2017), as their aggregating approaches are counting each value of all types’ ES (Costanza et al., 2014; Dai et al., 2016) (Figure 5). Stoeckl et al. (2014) try to consider the complexity and non-linearity of ecosystems and use statistical techniques to identify and control their overlapping benefits, but their methods and estimates still have some limitations. In DVS, the total value of ES in a region (or the Earth) takes the maximum one from the values of all types’ ES (Figure 3, 5). So, the severities of underestimate or overestimate on the overall ES value in a region (or the Earth) led by miss-counting or double-counting of ES types in ES valuation would be relieved in DVS, although DVS do not completely avoid the miss-counting or double-counting of ES types led by insufficient information (Figure 5).

**Figure 5.**
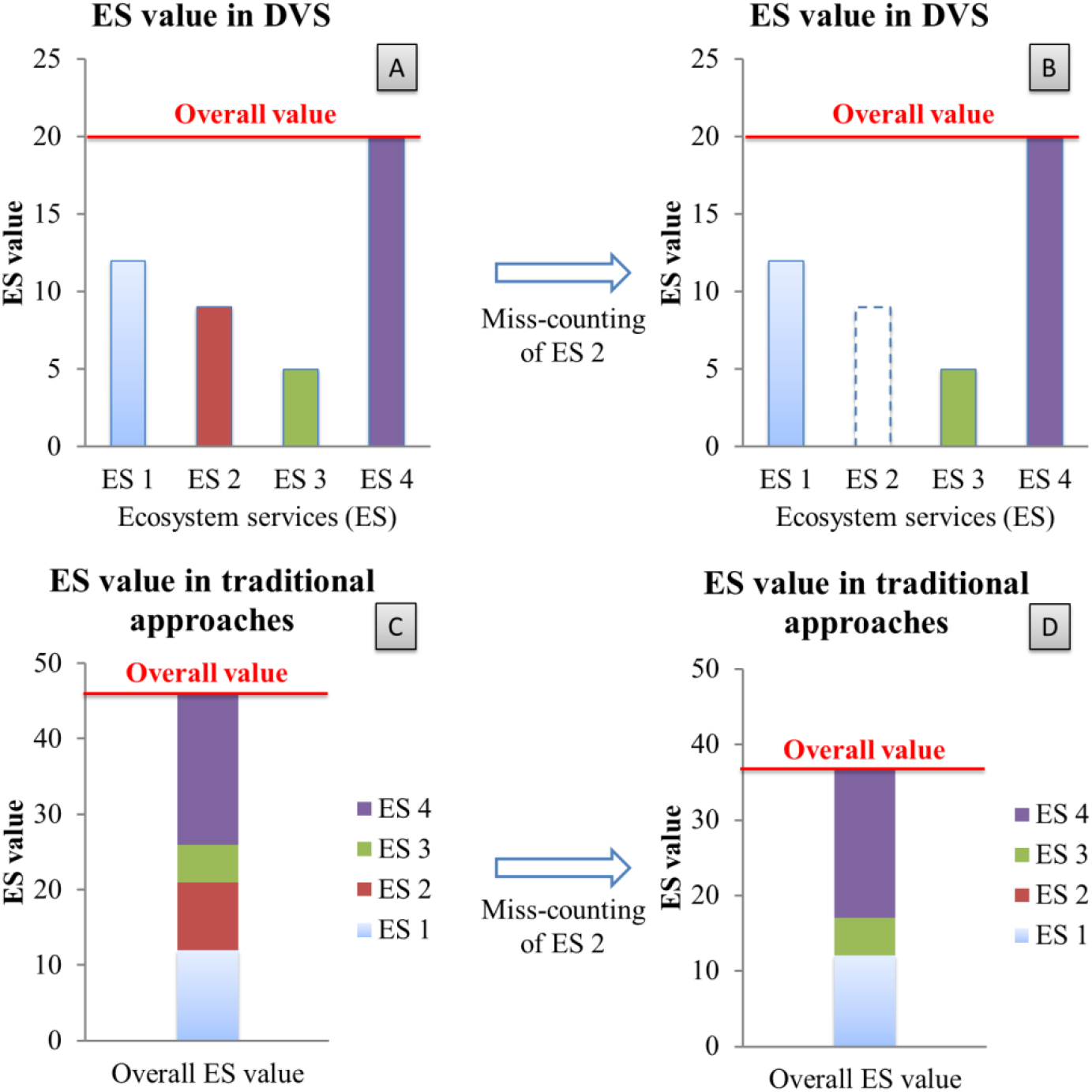
The schematic diagrams for the severities of underestimate on the overall ES value in a region (or the Earth) led by the miss-counting of ES 2 in two ES valuation approaches. The miss-counting of ES 2 does not lead a severe underestimate on the overall ES value in DVS (from A to B), but does in traditional approaches (from C to D). The values in these figures are all fictitious.

## 5. Conclusions

In this paper, we proposed a dual value system (DVS), a new assessment scheme to value ES. DVS includes two types of ES value: (1) Output Support Value (OSV) means the TVPH supported by the consumed ES, and (2) Optional Capacity Value (OCV) means the optional capacity of supporting the TVPH provided by the total ES. The overall value of ES in a region (or the Earth) takes the maximum one from the values of all types’ ES.

DVS provides new insight to understand and value ES:

1. DVS makes the importance and value of ES easy be understood in new perspective;
2. DVS concisely shows the relationship between ES value and the social and economic development;
3. DVS provides a tool for a new PES approach;
4. DVS provides a new framework for ES management;
5. DVS has the scale-specificity.

DVS overcomes the limitations on current ES valuation approaches:

1. DVS avoids the practical difficulties in traditional ES valuation approaches;
2. DVS makes the misestimate of ES value traceable and easy-to-correct;
3. DVS provides a general approach and a general unit to value ES;
4. DVS avoids the amplification of underestimate or overestimate on the overall ES value of a region.

DVS provides a useful quantifying framework for coordinating ecological protection and ecological compensation.

## Acknowledgments

We thank Li Yongchi, Li Zhaolei, Yang Yuanyuan and Zhao Bin for grateful assistance that helps to improve the manuscript. This research has not received any specific grant from funding agencies in the public, commercial, or not-for-profit sectors.

ES: ecosystem service(s)
OSV: output support value
OCV: optional capacity value
DVS: dual value system
GDP: gross domestic product
TVPH: total value produced by human being’s economic and social activities
PRWRC: Pearl River Water Resources Commission of the Ministry of Water Resources

